# Adrenal Gland Macrophage-derived TGF-β Governs Vascular Permeability to Drive Monocyte Recruitment during Stress

**DOI:** 10.64898/2026.05.18.726080

**Authors:** Yingzheng Xu, Michael T. Patterson, Michael D. Chang, Ian Ahlberg, Chi-Yun Liu, Charles Roll, Hannah Hillman, Ainsley Kennedy, Patricia R. Schrank, Gage M. Stuttgen, Yoshiaki Kubota, Stoyan Ivanov, Bryce A. Binstadt, Jesse W. Williams

## Abstract

The adrenal glands are central regulators of systemic stress responses through tightly controlled glucocorticoid production. Yet, the contribution of local immune–vascular interactions to adrenal stress adaptation remains poorly understood. Here, we investigated the role of adrenal gland macrophages in coordinating stress-induced immune remodeling and vascular function. By integrating single-cell RNA sequencing datasets across four distinct stress models, including acute cold exposure, chronic social defeat, chronic inflammation, and systemic *Candida albicans* infection, we identified a conserved increase in monocyte recruitment to the adrenal gland, accompanied by dynamic macrophage transitions. Comparative transcriptomic and ligand–receptor analyses identified transforming growth factor-β (TGFβ) as a dominant macrophage-derived signal targeting adrenal endothelial cells across all stress conditions. Pharmacological blockade of TGFβ receptor signaling reduced endothelial activation, vascular permeability, and monocyte infiltration into the adrenal gland following stress, without directly altering resident macrophage numbers. Using genetic fate-mapping and conditional knockout models, we demonstrate that macrophage-derived, but not endothelial-derived, TGFβ is required to promote enhanced endothelial adhesion molecule expression, vascular fenestration, permeability, and efficient monocyte recruitment. Loss of macrophage TGFβ production also led to exacerbated systemic stress hormone levels. Together, these findings uncover a previously unrecognized macrophage-endothelial axis in the adrenal gland, whereby macrophage-derived TGFβ regulates vascular properties to support immune cell recruitment and stress adaptation. This immune-vascular crosstalk provides new mechanistic insights into adrenal homeostasis and suggests potential therapeutic avenues for disorders associated with dysregulated chronic stress.

## Introduction

The adrenal glands are essential endocrine organs that regulate the body’s response to environmental and internal stressors by producing steroid hormones, including glucocorticoids and mineralocorticoids^1–4^. Glucocorticoid secretion is tightly controlled by the hypothalamus-pituitary-adrenal (HPA) axis, where stress-induced signaling promotes the release of corticotropin-releasing hormone (CRH) from the hypothalamus, adrenocorticotropic hormone (ACTH) from the anterior pituitary gland, and ultimately glucocorticoids from the adrenal cortex^5– 7^. Glucocorticoids are discretely released from spatially restricted zonation within the adrenal gland, which is critical for maintaining physiological homeostasis, modulating insulin sensitivity, vascular tone, and immune function^8–10^. However, sustained elevation in glucocorticoids has been implicated in the development of metabolic and cardiovascular diseases, including obesity and atherosclerosis, underscoring the importance of understanding the regulatory mechanisms governing adrenal hormone production^11–13^.

Macrophages, a prominent immune cell population found in all tissues, play pivotal roles in maintaining tissue homeostasis and immune regulation^14–16^. Tissue-resident macrophages exhibit marked diversity, with their phenotypes and functions shaped by the local microenvironment^17,18^. For example, macrophages in brown adipose tissue regulate thermogenesis^19–21^, while splenic red pulp macrophages capture iron for recycling^22–24^. In endocrine organs, macrophages contribute to tissue-specific functions, yet their roles in regulating hormone production remain poorly understood^25–27^. Recent work identified a distinct population of macrophages within the adrenal gland that exhibit sex-specific differences and are developmentally regulated by sex hormones^28–30^. Notably, adrenal macrophage depletion is associated with shifts in aldosterone production, even in animals lacking external stressors^28^. Further studies showed that following stress stimuli, adrenal gland macrophages adopt a lipid associated macrophage phenotype, supported by the infiltration of newly recruited monocytes^29,31,32^. Adrenal lipid associated macrophages can participate in stress hormone regulation through a macrophage lipid sensor, Trem2, which was demonstrated to control TGFβ secretion and limit corticosteroid production in adrenal cortical cells^29,31,33^. While, this work provides important insights into the endocrine-regulatory capacity of adrenal macrophages, the extent to which adrenal gland macrophages regulate other physiological properties of the adrenal gland remains unclear^34^.

In this study, we utilized single-cell RNA sequencing (scRNAseq) data to investigate conserved stress-regulatory mechanisms by which adrenal gland macrophages deploy across several different stimuli. We incorporated sterile cold stress, sterile chronic inflammation^29^, chronic social stress^35^ and *Candida albicans* infection^36^ to explore transcriptional models of cell-cell interaction within the adrenal gland. These studies highlighted the potential role of TGFβ in coordinating endothelial activity and response to stress. Macrophage specific TGFβ deletion using an in vivo mouse model revealed that TGFβ was required to modulate endothelial permeability, which directly impacted monocyte recruitment to the adrenal gland under stress conditions. Thus, reducing the ability of monocyte-derived lipid associated macrophages to form within the adrenal gland, which led to exacerbated stress responses. This study highlights the significance of cellular crosstalk between macrophages and the vascular system within the adrenal gland and offers potential therapeutic insights for future investigations into settings of chronic hypercortisolism.

## Results

### scRNAseq integration of the adrenal gland under different stress stimulation

To define how adrenal gland immune populations respond to diverse forms of stress, we performed an integrated scRNAseq analysis of adrenal gland cells collected from mice exposed to 4 distinct stress stimuli including acute cold exposure, chronic social defeat stress, acute systemic *Candida albicans* infection, and chronic atherosclerosis^29,35,36^ (Fig 1A). A total of 44,306 cells passed quality control and were included in integration and downstream analysis. 17 clusters were identified, followed by an unbiased cell annotation approach performed using SingleR^37^ (Fig 1B). Notably, a robust macrophage population, cluster 1, was revealed in all stress conditions, despite a lower frequency of these cells in the chronic social stress dataset (Fig 1C). To profile genes representing each cluster, we performed a differential analysis among all clusters and highlighted the top 3 genes that defined that cluster (Fig 1D). Clusters 0, 1, 11, 14, and 16 are dominated by macrophage genes featuring *C1qa, C1qc, and Apoe*, with cluster 11 showing high proliferative capacity indicated by high *Mki67* expression. Clusters 2, 3, 5, 10, and 13 exhibited high *Cyp11b1, Cyp21a1, and Star* expression, suggesting that these cells are primarily adrenal endocrine cells. Cluster 4 was predominantly monocytes indicated by the expression of *S100a4 and Plac8*. Cluster 6 consisted mainly of B cells with high *Igkc and Iglc2*, suggesting immunoglobulin production. Interestingly, cluster 14 which is a macrophage cluster, also contains several B cell associated genes, suggesting this cluster may also contain B cell precursors (pro-B cells) or transcripts detected in macrophages that have engulfed B cells. Cluster 8 showed significant *Cd3d and Cd3g* expression, therefore suggesting a T cell population. *Nkg7, Ccl5, and Gzma* identified natural killer cells in cluster 9. Cluster 12 features *S100a9 and S100a8*, consistent with a neutrophil origin. High *Scg2, Chgb, and Chga* expression indicates another endocrine cell population in cluster 15, likely neuroendocrine cells. Of the immune compartment, the social stress study was primarily composed of macrophages, but had very low immune cell diversity, which was consistent with a control dataset obtained at the same time. All immune categories could be found in cold stress, atherosclerosis, and acute *Candida* infection datasets (Fig 1E). Next, we examined the immune dynamic changes within each stress condition. Interestingly, we observed a drastic increase in the expression of CCR2+ cell, indicating monocytes and monocyte-derived macrophages, shown in red on pie charts, in all stress settings except social stress (Fig 1F). The lack of response in the chronic social stress model may be a result of the sustained glucocorticoid exposure in this model^38,39^. Together, these data suggest that monocyte recruitment is accelerated in the adrenal gland in response to diverse stress stimuli.

**Fig 1.**
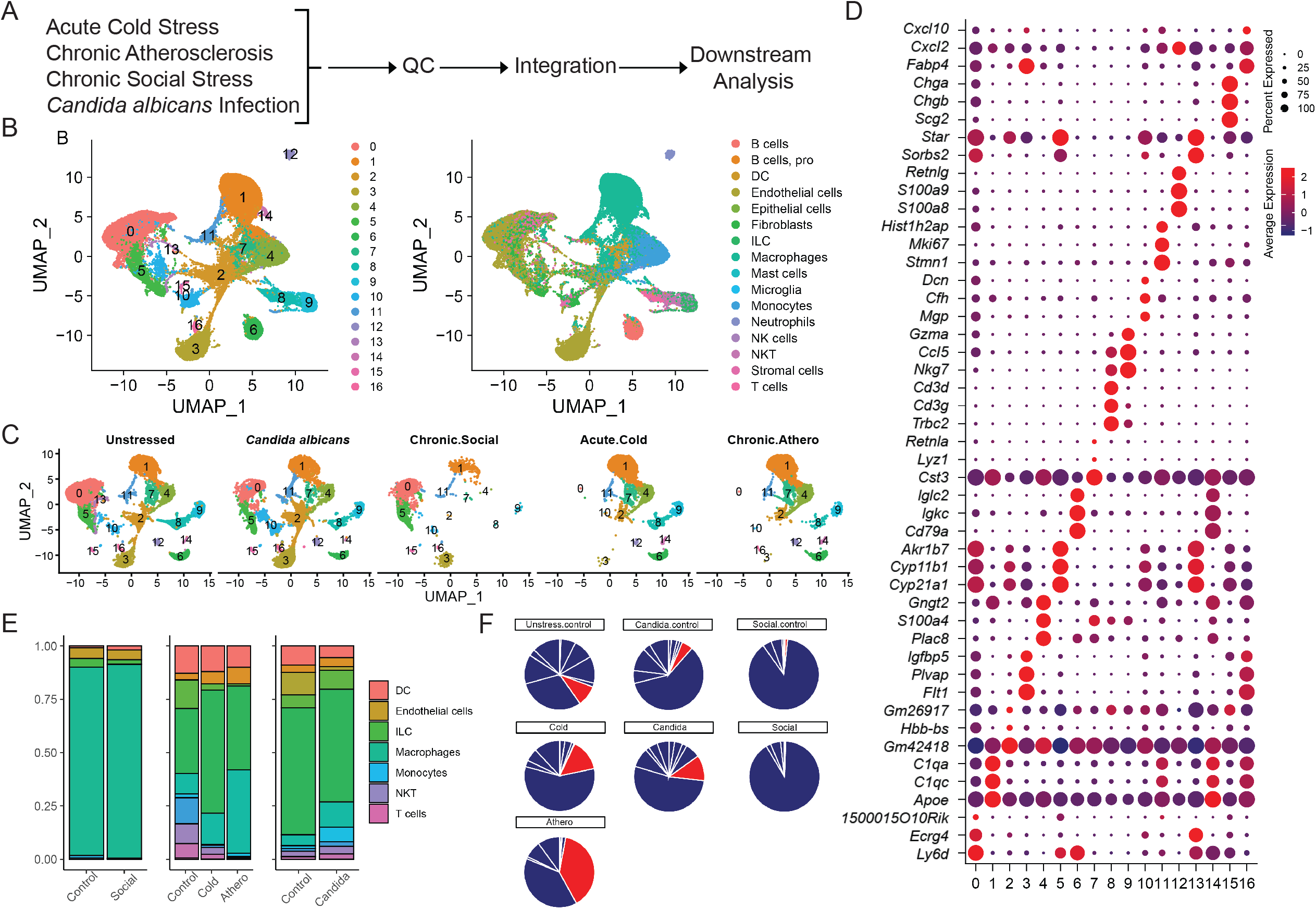
scRNA-seq integration of adrenal gland stress datasets. A. scRNAseq integration of sterile and non-sterile stress conditions, including basal unstressed, acute cold, chronic inflammation, chronic social defeat, and acute *Candida albicans* infection. B. UMAP showing clusters and cell identities in the integrated analysis. C. UMAP clusters split by stress condition. D. Top three differentially expressed genes for each cluster. E. Immune composition across stress conditions. F. CCR2+ cell proportion, indicating monocytes and monocyte-derived macrophages, for each stress condition (shown in red).

Next, we sought to validate this observation using mouse models of cold exposure as a reproducible approach for inducing acute stress. We exposed C57BL/6 (B6) mice to 4°C for 24 hours to induce cold stimulation and quantified adrenal gland monocytes using flow cytometry (Fig 2A, Supplemental Fig 1A). In line with our computational observation, Ly6C+ classical monocytes accumulated significantly more in cold-stressed mice compared to unstressed mice (Fig 2B, Fig 2C). This result is consistent with prior work showing that recruited monocytes replaced adrenal macrophages following stress^29^. To further explore this observation, we utilized a genetic monocyte fate mapping approach using a *Ccr2*^*creER*^ *R26*^*TdTomato*^ reporter mouse strain. This approach specifically labels classical monocytes in the blood^40^, allowing for tracking the recruitment and differentiation of monocytes into macrophages with expression of the tdTomato fluorescent protein. We pulsed mice with oral tamoxifen administration 6 or 24 hours before sacrifice to label CCR2-expressing cells and map their recruitment in early and late waves of infiltration to the adrenal gland during cold challenge (Fig 2D). Interestingly, we observed a cluster of monocytes recruited to the adrenal cortex and surrounding the adrenal medulla at 6 hours post tamoxifen stimulation, where a small number of monocytes had already started differentiating into macrophages. At 24 hours, monocytes were evenly distributed in the adrenal gland, and more cells initiated differentiation, suggesting that monocytes enter the adrenal gland from the vasculature surrounding the medulla (Fig 2E). Additionally, we were curious about the degree to which circulating monocytes impact the monocyte population in the adrenal gland. We utilized *Ccr2*-deficient mice, possessing a GFP knock-in allele (*Ccr2*^*GFP/GFP*^), as these mice show reduced circulating monocytes in blood (Fig 2F, Supplemental Fig 1A). As expected, we found reduced classical monocytes in *Ccr2*-deficient mice in both circulation (Supplemental Fig 1B-D) and the adrenal gland following stress^41^ (Fig 2G-I). Together, these data suggest that stress promotes CCR2-dependent monocyte infiltration to the adrenal gland, which supports local macrophage population maintenance.

**Fig 2.**
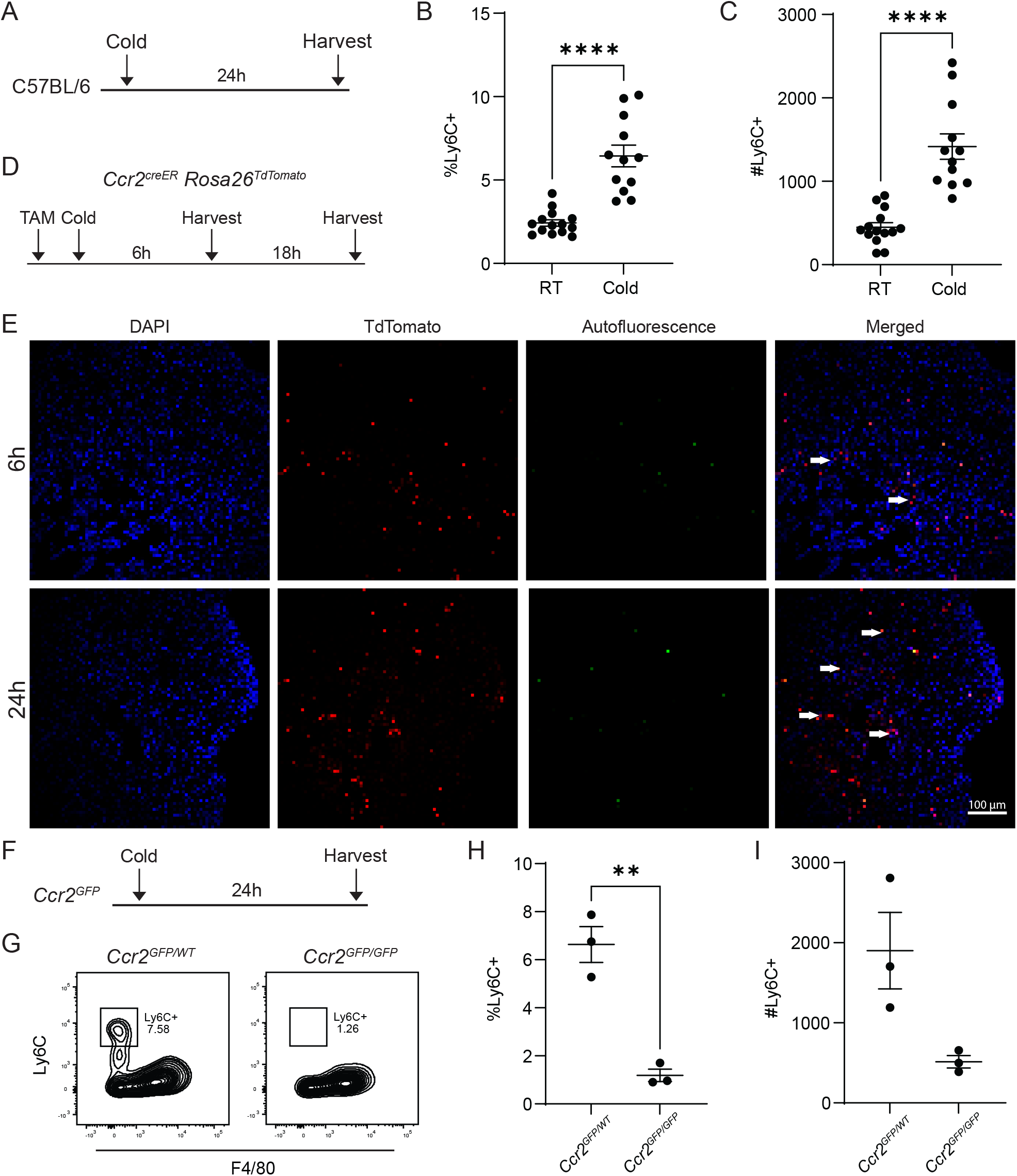
Fate mapping shows enhanced monocyte entry and differentiation in adrenal gland following stress. A. Experimental setup for the acute cold stress model. B6 mice were cold-stressed in 4°C for 24 hours. B, C. Quantification of Ly6C+ classical monocyte in unstressed mice at room temperature (RT) or cold-stress mice adrenal glands. D. Experimental setup for monocyte fate mapping under cold stress. E. Localization of TdTomato+ cells in the adrenal gland post 6 and 24 hours of TAM administration. F. Experimental setup for CCR2 deletion experiment. G. Representative flow cytometry plot showing efficient deletion by homozygous CCR2 GFP/GFP. H-I. Quantification of Ly6C+ classical monocytes in CCR2 sufficient and deficient mice adrenal glands.

### Investigation of the shared mechanisms underlying adrenal stress stimulation

Our prior work highlighted the heterogeneity of adrenal macrophages and monocytes following acute and chronic physiologic stressors^28,29^, where we documented a lipid-associated macrophage population that plays vital roles in adrenal stress hormone homeostasis. Thus, we investigated the subtypes of adrenal gland macrophages under acute *Candida* infection, chronic social defeat stress, cold stress, and atherosclerosis. We first re-clustered the monocytes and macrophages to explore potential dynamic changes of macrophage subtypes associated with stress stimulations. Re-clustering revealed 9 subpopulations (Fig 3A) where 0, 1, 3, 4, 5, 7, and 8 were identified as macrophages. Notably, all subclusters were equally composed of cells from different stress conditions, except clusters 2, 3, and 6, which had a higher proportion of cells from atherosclerosis challenge. Similarly, *C. albicans* infection was found to dominate cluster 5 (Fig 3B). We also explored the cluster composition of each stress condition and found a balanced distribution of clusters in all conditions, with an exception that social stress largely composed of macrophage clusters 0 and 1 (Fig 3C-D). Next, we explored the transcriptional features of these subpopulations and stress conditions to examine transcriptional similarities and differences underlying different stress challenges. First, we looked at the top 5 features leading each subcluster (Fig 3E). Clusters 2, 3, and 6 showed high expression of *Hp, Ccr2, S100a10*, and *Ly6c2*, suggesting a monocyte population. Clusters 0 and 5 are identified as chemokine-producing macrophages, as these cells feature *Ccl4, Ccl3, Cxcl2, and Cxcl1*, with the addition that cluster 5 showed prominent proliferative capacity reflected by *Mki67*. Clusters 1 and 4 exhibited consistent *C1qb and C1qa* expression, suggesting a macrophage lineage. Cluster 4 showed relatively high expression of *Folr2* that likely suggests an alternatively activated macrophage population^17,42–46^. Although clusters 7 and 8 exhibit macrophage features, these populations also show B cell signatures, such as *Ly6d, Cd79a, Gzma, and Ms4a4b*, suggesting a contaminated population (Fig 3E).

**Fig 3.**
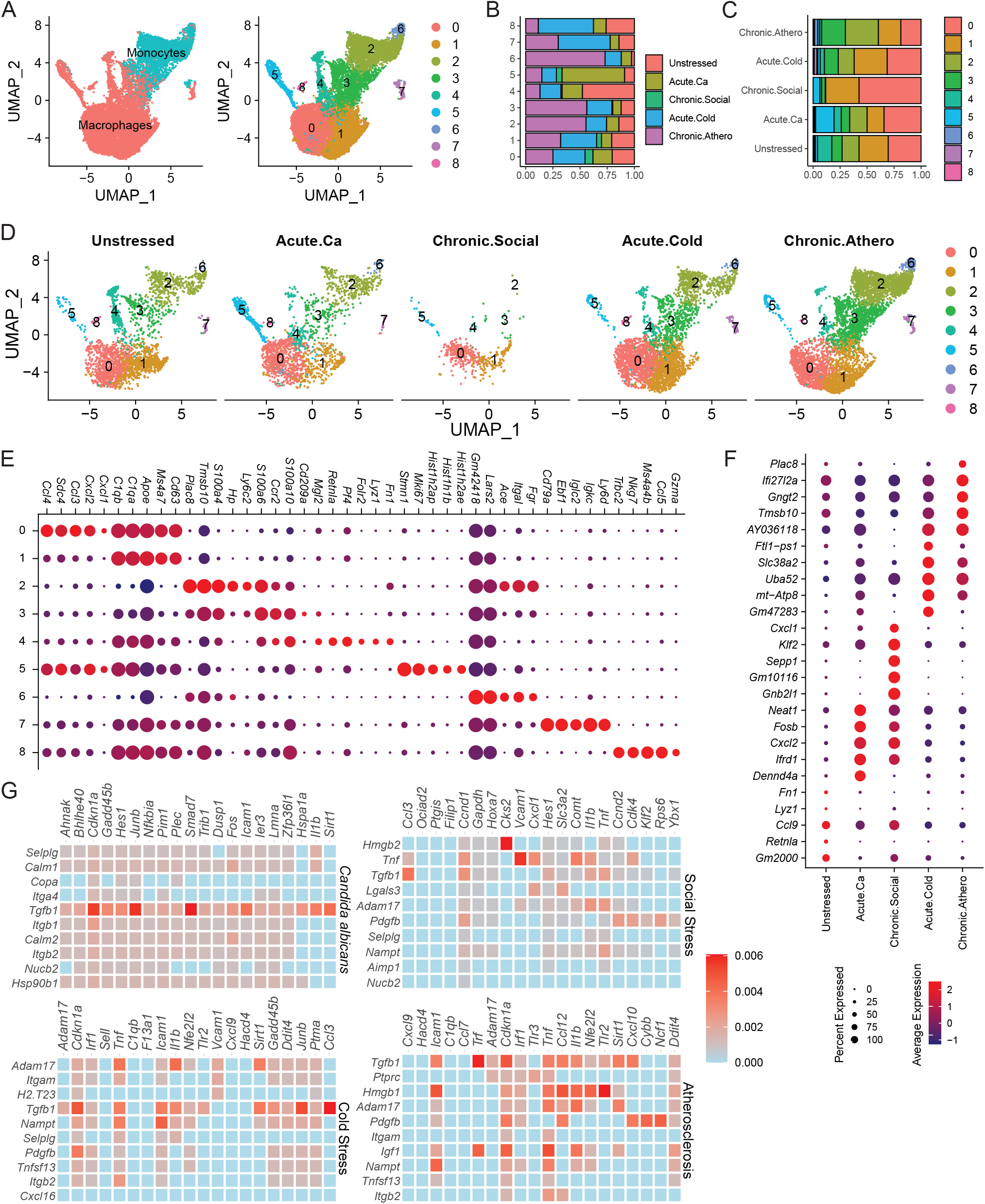
Re-clustering of adrenal gland macrophages and cell communication with endothelial cells. A. Re-clustering of monocytes and macrophages displayed in UMAP space. B–C. Cluster and condition composition for the UMAP clusters shown in A. D. UMAP clusters split by stress condition. E. Representative differentially expressed genes (DEGs) for each cluster. F. Representative DEGs for each condition. G. NicheNet analysis showing macrophage– endothelial cell crosstalk under different stress conditions.

Furthermore, we asked whether macrophages undergo unique transcriptional alterations in response to different stress stimuli. To investigate this idea, we performed differential expression analysis comparing stress conditions of the monocyte and macrophage populations (Fig 3F). Compared to unstressed mice, acute *C. albicans* infection led to high expression of *Neat1, Fosb, Cxcl2, Ifrd1, and Dennd4a*, suggesting a response to inflammation, immune regulation, and cellular stress^47–52^. Interestingly, a non-pathological stress like social defeat also resembled these features, with an additional upregulation of *Gnb2l1, Sepp1, Klf2, and Cxcl1*, which also suggested a mixed stage of inflammation^53–56^. Similar to prior reports, cold stress and atherosclerotic stress exhibited overlapping transcriptional features^29,57^, as these two conditions both upregulate *Plac8, Ifi27l2a, Gngt2, Tmsb10, and Slc38a2*, which likely represent a complex macrophage transitioning stage involving responses to immune modulation, metabolic reprogramming and macrophage polarization^58–62^. To contextualize the stress-dependent transcriptional states of macrophages within the broader tissue environment, we next explored macrophage regulation of endothelial activation. According to a recent report emphasizing the role of macrophages in adrenal endothelial cell (EC) specialization^63^, we hypothesized that adrenal macrophages may contribute to stress regulation by modulating functions of endothelial cells. To investigate potential signaling communicated between EC and macrophages, we applied NicheNet package in R to predict ligand-target interactions. Across all stress settings, we observed a strong outgoing signal of *Tgfb1* from macrophages that impacts various EC gene signatures (Fig 3G). Notably, TGFβ was the only macrophage-secreted molecule that elicited a strong signal to engage ECs across all 4 stress conditions. In conclusion, these data suggest that macrophages may regulate EC function via TGFβ to modulate stress response in adrenal gland.

### TGFβ promotes adrenal gland monocyte recruitment

To examine the impact of TGFβ signaling on adrenal gland ECs, we blocked TGFβ receptor (TGFβR) using Ly573636, a small molecule TGFβRI antagonist, in B6 mice in vivo (Fig 4A). First, we used flow cytometry to quantify adrenal gland macrophages in Ly573636 or vehicle-treated mice following cold exposure or maintained at room temperature (Supplemental Fig 2A). No difference was observed associated with cold stress or TGFβR signaling blockage (Supplemental Fig 2B-C), which suggests that TGFβ signaling did not directly influence the mature macrophage population. We then switched attention to the adrenal gland monocyte population, as these were dramatically shifted in the scRNA-seq analysis. After cold stress, vehicle-treated mice showed a significant increase in the recruitment of local adrenal gland monocytes, which was completely attenuated in Ly573636 treatment (Fig 4B-D). Next, we hypothesized that TGFβ influenced EC adhesion molecule expression that regulates the rate of monocyte influx. Therefore, we measured Vcam1 protein expression among adrenal gland ECs. As expected, cold stress promoted Vcam1 expression on adrenal gland ECs, whereas TGFβR signaling blockage reversed this induction (Fig 4E-F, Supplemental Fig 2A). This phenotype suggested TGFβ-driven EC activation, likely directly linked to monocyte recruitment and infiltration. Consistent with this concept, we tested whether cold exposure affected vascular permeability in the cold-exposure model using Evans blue dye (EBD) injection in vivo. Notably, cold-stressed mice showed a greater degree of vascular permeability, which was inhibited by TGFβR blockade (Fig 4G-H).

**Fig 4.**
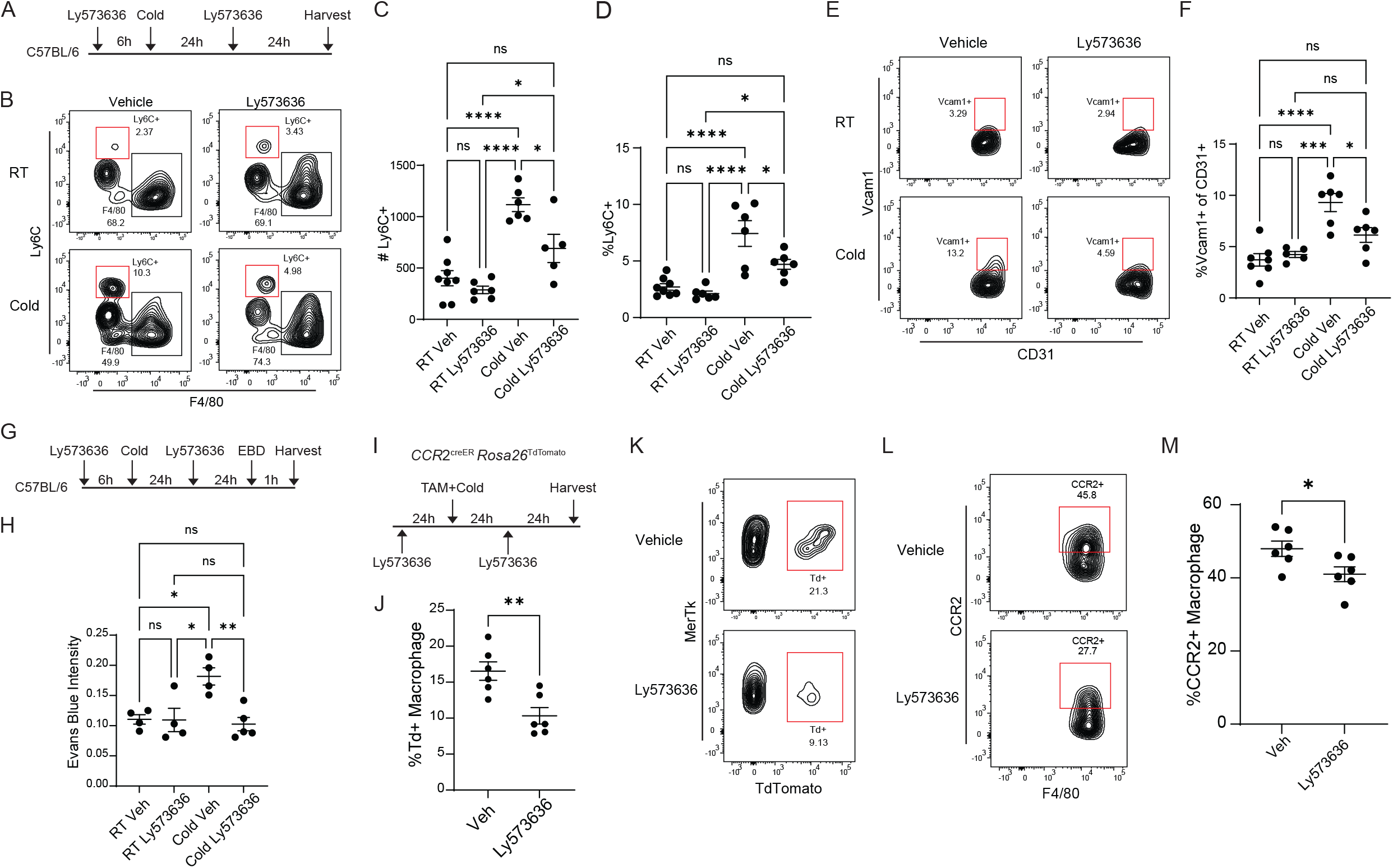
Therapeutic blockade of TGFβR reduces monocytes in the adrenal gland. A. Experimental setup for blocking TGFβR signaling in cold-stressed mice. B. Representative flow cytometry plot showing reduced classical monocytes in the adrenal gland after TGFβR blockade. C–D. Quantification of classical monocytes. E. Representative flow cytometry plot showing vascular cell adhesion molecule 1 (VCAM1) expression on (CD31+) endothelial cells. F. Quantification of endothelial VCAM1 in unstressed or cold-stressed mice receiving vehicle control or TGFβR blockade. G–H. Evans blue dye assay experimental design, representative flo cytometry plots, and quantification showing endothelial permeability in mice with or without TGFβR blockade. I–K. Monocyte fate mapping experimental setup and quantification in mice with or without TGFβ receptor blockade. L–M. Representative flow cytometry plots and quantification of CCR2^+^ macrophages in mice with or without TGFβ receptor blockade.

Next, we were curious whether TGFβ also impacted monocyte differentiation to adrenal macrophages. We designed a monocyte fate mapping experiment to investigate this possibility, where *Ccr2*^*creER*^ *R26*^*TdTomato*^ mice were used to quantify the percentage of macrophages that are turned over from newly recruited cells under TGFβR antagonism in stress (Fig 4I). Mice were first given 1 dose of Ly573636 to block TGFβ signaling, followed by TAM administration to activate tdTomato expression in *Ccr2*-expressing cells and cold housing 24h after the initial Ly573636 treatment. Mice were maintained in the cold for 48h and received another Ly573636 treatment during cold incubation. By extending this analysis window beyond the 24 hours that we previously explored, we could detect substantial tdTomato+ macrophage accumulation within the adrenal gland. Interestingly, Ly573636-treated mice showed a smaller tdTomato+ macrophage ratio, suggesting a slowed monocyte recruitment rate in these mice compared to vehicle treated control mice (Supplemental Fig 3A, Fig 4J-K). Notably, the tdTomato expression induced by *Ccr2*^*creER*^ *R26*^*TdTomato*^ is highly specific to the monocyte lineage, where only classical and nonclassical monocytes (which mature from CCR2+ classical monocytes) showed detectable tdTomato signal in peripheral blood (Supplemental Fig 3B-L). We also examined the accumulation of CCR2+ monocyte-derived macrophages, as these are potential transitioning macrophages from monocytes. Notably, Ly573636-treated mice exhibited fewer CCR2-expressing macrophages (Fig 4L-M). Together, we utilized Ly573636 and an *in vivo* system to demonstrate that TGFβR signaling modulates adrenal gland vascular permeability, monocyte recruitment, and their subsequent differentiation into adrenal macrophages.

### Macrophage TGFβ regulates vascular permeability and monocyte recruitment following cold stress challenge

TGFβ is produced from multiple sources and found in all tissues in mice^64,65^. In the adrenal gland, *Tgfb1* is readily detected in both macrophages and ECs, indicated by high *C1qc* and *Pecam1* expression, respectively (Supplemental Fig 4). Based on our cell-cell communication analysis using NicheNet, we first examined the effects of myeloid cell-specific TGFβ deletion in macrophages and its effect on adrenal EC function modulation in our model of stress, using the *Cx3cr1*^*creER*^ *Tgfbβ1*^*flox/flox*^ (*Tgfb1*^ΔMac^) mice (Fig 5A). CX3CR1 is highly expressed by adrenal macrophages and circulating monocytes, as well as some dendritic cells. Importantly, *Cx3cr1* is not robustly expressed by ECs. Thus, it is an efficient model to test the impact of myeloid specific deletion of *Tgfb1*. To ensure this cre-flox system efficiently deleted TGFβ, we used confocal microscopy to visualize cells that express TGFβ in *Tgfb1*^ΔMac^ or WT mice. *Tgfb1*^ΔMac^ prominently lost TGFβ in the adrenal gland cortex when compared to control littermates following cold challenge (Supplemental Fig 4A). These data also showed a nearly complete loss of TGFβ in the adrenal parenchyma, suggesting that macrophages may be the primary source of TGFβ in adrenal glands. SMAD activation and signaling is the primary signaling pathway downstream of TGFβR signaling ^66–70^. Therefore, we proposed that, in the absence of TGFβ, SMAD signaling will likely be reduced in*Tgfb1*^ΔMac^ mice. We performed confocal microscopy to test this possibility. As expected, phosphorylated SMAD2 was significantly reduced in *Tgfb1*^ΔMac^ mice compared to controls under acute cold stress settings, showing that the signaling downstream of TGFβR was also attenuated when TGFβ was deleted in macrophage lineage cells (Supplemental Fig 4B).

**Fig 5.**
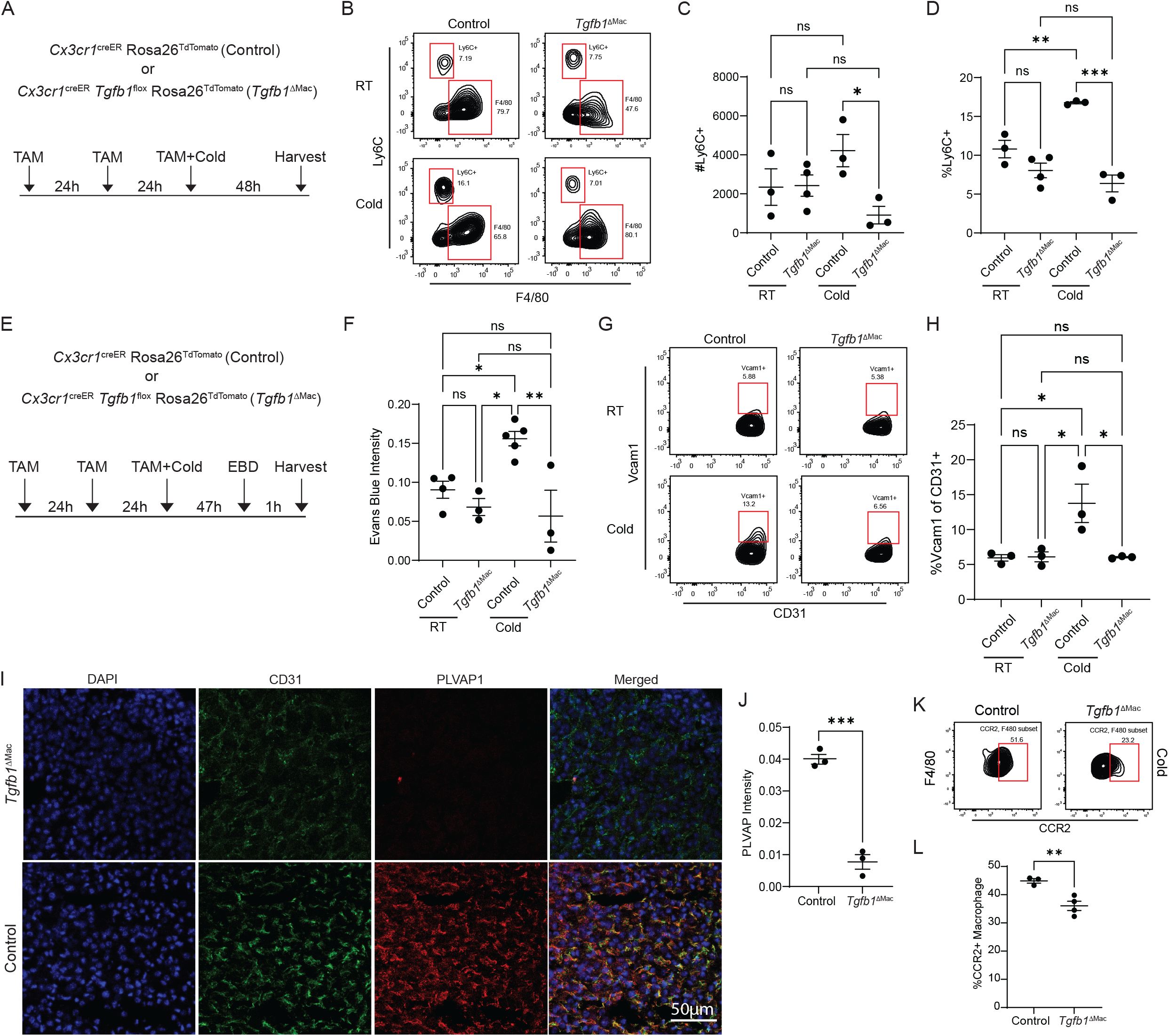
Macrophage-specific TGFβ1 deletion leads to reduced adrenal gland monocytes and vascular permeability. A. TGFβ1 deletion under cold stress using *Cx3cr1*^creER^ *Tgfβ1*^flox^ system experimental setup. B– D. Quantification of Ly6C^+^ monocytes in adrenal glands from TGFβ1-deficient or control mice. E. Experimental setup of vascular permeability using Evans blue dye (EBD) in TGFβ1-deficient mice. F. Quantification of EBD intensity in TGFβ1-deficient mice or controls. G–H. Representative flow cytometry plots and quantification of VCAM1 in adrenal gland endothelial cells from TGFβ1-deficient mice or controls. I. Confocal microscopy micrographs of adrenal gland vascular fenestration measured by PLVAP1 staining (red). J. Quantification of fluorescence intensity of PLVAP1 staining. K–L. Representative flow cytometry plots and quantification of CCR2^+^ macrophages in TGFβ1-deficient mice or controls.

In *Tgfb1*^ΔMac^ mice, we first investigated the monocytes in the adrenal gland. Similar to the phenotype induced by TGFβR blockade, macrophage-specific TGFβ deletion also significantly reduced Ly6C+ monocyte recruitment to the adrenal gland following cold stimulation (Fig 5B-D), suggesting the necessity of macrophage-derived TGFβ in vascular function. Next, we tested vascular permeability with an Evans blue dye experiment in *Tgfb1*^ΔMac^ mice. Consistent with the reduced monocyte counts, *Tgfb1*^ΔMac^ mice also showed reduced vascular leakiness in the adrenal gland, confirming that macrophage-derived TGFβ mediates EC responses to cold stress (Fig 5E-F). Furthermore, we found that Vcam1 was greatly reduced in *Tgfb1*^ΔMac^ mice compared to control mice following stress challenge, once again suggesting the importance of TGFβ in EC function during stress (Fig 5G-H). Additionally, increased vascular permeability is often related to higher fenestration of the vessel^71–73^. Therefore, we hypothesized that *Tgfb1*^ΔMac^ mice may exhibit reduced fenestration due to TGFβ deletion, which was reflected as reduced vessel permeability. To test this, we utilized confocal microscopy to measure plasmalemmal vesicle-associated protein (PLVAP1), because it supports the fenestrae in fenestrated vessels^63,74,75^. In line with our observation, PLVAP1 was drastically lower in *Tgfb1*^ΔMac^ mice compared to controls (Fig 5I-J), suggesting reduced vessel permeability is also associated with reduced EC fenestration. Strikingly, we found that not only fenestration, but the overall EC morphology was altered, showing reduced CD31 expression and potentially reduced density (Supplemental Fig 5A-B). Next, with respect to macrophage turnover, we examined the CCR2+ macrophage population induced by stress in the control and *Tgfb1*^ΔMac^ mice. CCR2-expressing transitioning macrophages were reduced in the absence of macrophage-produced TGFβ (Fig 5K-L). Moreover, we found higher corticosterone levels in the serum of *Tgfb1*^ΔMac^ mice, which is consistent with our prior study that supported a role for monocyte-derived lipid associated macrophages in the regulation of steroidogenesis in the adrenal gland^29^ (Supplemental Fig 6A). With this evidence showing the impact of macrophage TGFβ on ECs, we concluded that macrophage-derived TGFβ was vital for adrenal gland EC maintenance.

**Fig 6.**
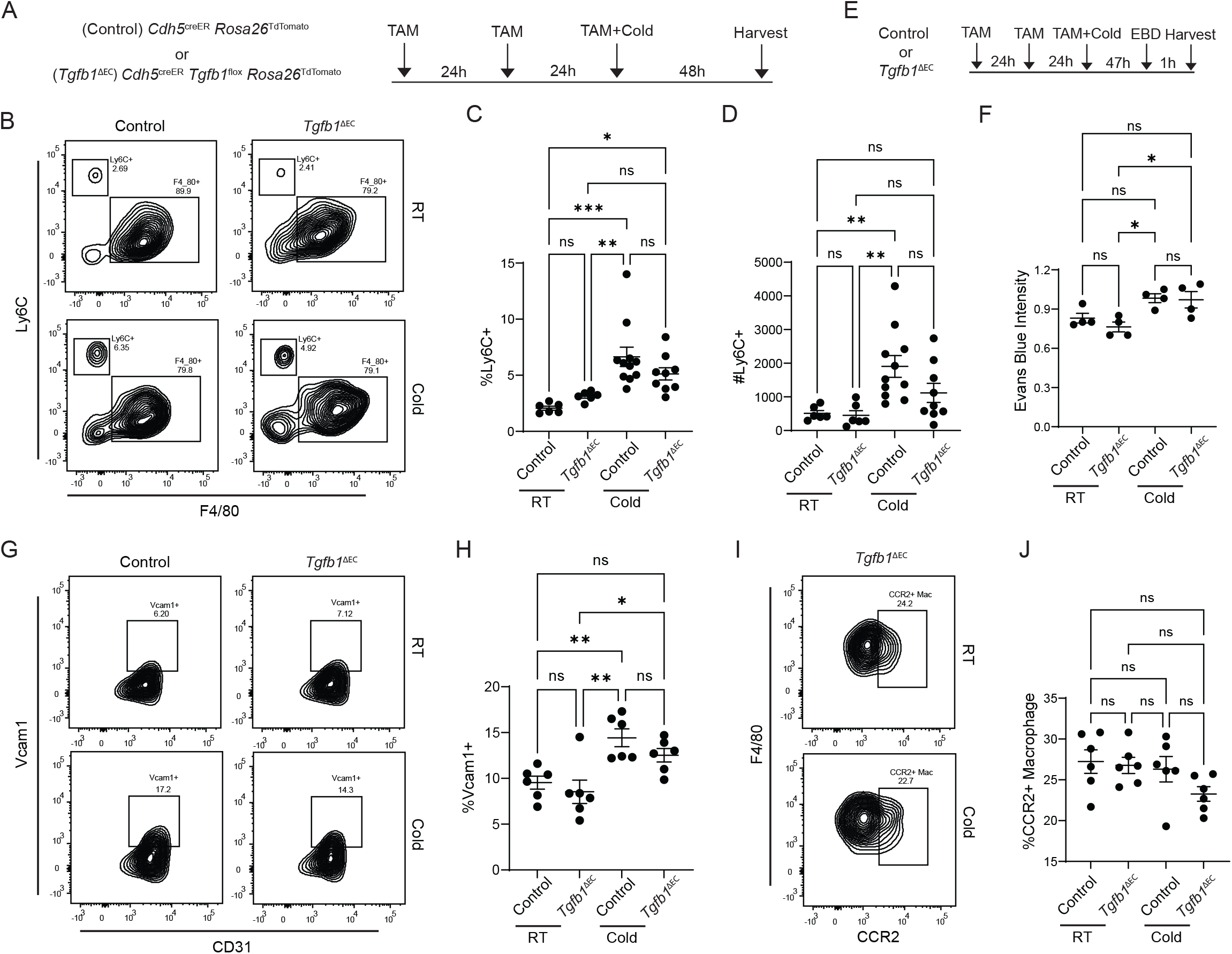
Endothelial TGFβ1 deletion has minimal effect on adrenal gland monocyte entry. A. Experimental setup of endothelial cell–specific TGFβ1 deletion using *Cdh5*^*creER*^ TGFβ1 flox system. B–D. Representative flow cytometry plots and quantification of Ly6C^+^ monocytes in adrenal glands from endothelial cell TGFβ1-deficient or control mice. E–F. Experimental setup and quantification of vascular permeability measurement using Evans blue dye in *Cdh5*^*creER*^ TGFβ1 flox mice. G–H. Quantification of VCAM1 in adrenal gland endothelial cells from endothelial TGFβ1-deficient or control mice. I–J. Representative flow cytometry plots and quantification of CCR2^+^ macrophages from endothelial TGFβ1-deficient or control mice.

Given that *Tgfb1* is also expressed by ECs, we next aimed to assess the impact of EC-originated TGFβ on monocyte recruitment and EC properties. For this purpose, we generated an inducible EC-specific TGFβ deletion model using *Cdh5*^*creER*^ *Tgfb1*^*flox/flox*^ (*Tgfb1*^ΔEC^) to investigate the effects of EC-derived TGFβ (Fig 6A)^76–79^. Following the same treatment protocol as outlined in the macrophage-specific deletion model, we examined the classical monocytes in the adrenal gland after TGFβ deletion in ECs. Compared to the WT setting, *Tgfb1*^ΔEC^ did not show a dramatic shift in the accumulation of Ly6C+ monocytes following cold stress, as both the proportion and absolute count of these cells did not result in statistically significant change (Fig 6B-6D). However, it is noted that there are minor trends observed that could suggest that EC-derived TGFβ might have a minor effect on the recruitment of monocytes. We also measured vascular permeability using Evans blue dye assay (Fig 6E). Consistent with the prior conclusion, there was no change in the permeability of the vasculature in the adrenal gland associated with EC-specific TGFβ deletion (Fig 6F). In addition, we measured the activity of Vcam1 to determine if TGFβ deletion impacted monocyte recruitment via adhesion mechanisms. In line with previous observation, no significant difference was observed between WT and *Tgfb1*^ΔEC^ mice, although as observed in the monocyte recruitment assay there may be a potential partial impact that did not reach statistical significance (Fig 6H). Macrophage turnover was also assessed in *Tgfb1*^ΔEC^ mice by quantification of CCR2+ macrophages in the adrenal gland. Again, no significance was observed between WT or TGFβ deficient settings (Fig 6I-6J). Together, these observations suggest that unlike macrophage-specific deletion, EC-derived TGFβ is not required for EC response to adrenal gland stress that mediates the recruitment of monocytes from blood.

## Discussion

This study identifies adrenal gland macrophages as active regulators of endothelial barrier properties and monocyte recruitment during stress adaptation. We demonstrate that adrenal macrophage-derived TGFβ signaling modifies endothelial adhesion molecule expression and vascular permeability, thereby facilitating monocyte entry into the adrenal tissue. These findings reveal an immune-vascular axis in the adrenal gland that supports macrophage turnover and immune remodeling under conditions of physiological stress, which has direct effects on the regulation of systemic glucocorticoid release.

Previous work has largely focused on neuro-endocrine regulation of the HPA axis, with comparatively little attention paid to the contribution of local adrenal immune cells^80–82^. While macrophages in other tissues have been implicated in vascular remodeling, for example, microglia shaping the blood-brain barrier or liver macrophages guiding sinusoidal organization, our results provide the first mechanistic evidence that adrenal macrophages orchestrate adrenal gland vascular function^83–85^. By regulating endothelial junction integrity and adhesion molecule expression, adrenal macrophages establish a permissive environment for monocyte infiltration, ensuring a dynamic macrophage population that adapts to stress-induced demands. In fact, a recent study has demonstrated that adrenal gland macrophages can fine tune aldosterone production and blood pressure by preserving endothelial cell specification^63^. Importantly, this work highlighted the overlap with vascular remodeling in human adrenal glands with overproduction of aldosterone^63^. This is direct evidence that adrenal macrophages can indeed modulate vascular properties that impact adrenal steroidogenesis.

Our work also highlights TGFβ as a central mediator of immune-endothelial communication in the adrenal gland. While TGFβ has well-established roles in vascular biology, fibrosis, and immune regulation, its function in endocrine organ vascular dynamics have remained largely unexplored. The finding that adrenal macrophage-derived TGFβ can alter endothelial permeability and adhesion suggests that macrophage-derived cytokines may provide a fine-tuning mechanism for organ-specific vascular responses. This has important implications for disorders of adrenal function, where altered vascular permeability and immune infiltration are hallmarks of disease.

There are several broader implications of these findings. First, they suggest that stress-induced monocyte recruitment to the adrenal gland is not a passive process driven solely by systemic cues but rather an active, macrophage-dependent program. Second, the discovery of adrenal macrophage-endothelial crosstalk raises the possibility that similar immune-vascular mechanisms exist in other endocrine tissues, such as the pituitary, thyroid, or pancreas. Third, our results may provide a mechanistic link between acute stress, adrenal inflammation, and the pathogenesis of chronic stress-related disorders, including Cushing’s syndrome, adrenal hyperplasia, and metabolic disease. Several questions remain to be addressed regarding how adrenal macrophage-derived TGFβ production is regulated under basal versus stress conditions, whether distinct adrenal macrophage subsets differentially contribute to endothelial modulation, and whether there are long-term consequences of enhanced monocyte recruitment for adrenal homeostasis and function. Future studies employing conditional genetic models, fate-mapping studies, and intravital imaging will be critical to dissect these mechanisms further. Additionally, extending these observations to human adrenal tissues, particularly in patients with hypercortisolism or adrenal tumors, will be essential to establish translational relevance.

In conclusion, this study establishes adrenal macrophages as key regulators of vascular function and immune cell recruitment to the adrenal gland during stress adaptation. By uncovering a previously unrecognized adrenal macrophage-endothelial axis, our work adds a new dimension to the understanding of immune-endocrine crosstalk and highlights the dynamic role of macrophages in shaping endocrine tissue function. These findings open new avenues for exploring macrophage-targeted therapies in adrenal disorders and stress-related disease.

## Methods

### Animal use

Mouse models involved in this study were acquired and crossed from C57BL/6J (Jax 000664), B6.129P2(C)-*Cx3cr1*^*tm2*.*1(cre/ERT)Jung*^ /J(Jax 020940), *Cdh5*^*creER*^ (Tg(Cdh5-cre/ERT2)#Ykub)^79^, *Tgfb1* ^*tm2*.*1Doe*^ /J (Jax 010721), B6.Cg-*Gt(ROSA)26Sor*^*tm9(CAG-tdTomato)Hze*^ /J (Jax 007909) and B6(C)-*Ccr2*^*tm1*.*1Cln*^/J (Jax 027619). All research animals were housed at the Research Animal Resources (RAR) facilities of the University of Minnesota, under specific pathogen-free conditions. Both male and female mice were used in equal numbers in all experiments. Mice had access to water and food ad libitum and lived in 12 hour light, 12 hour dark cycles. All experiments and procedures were approved by the University of Minnesota Medical School Institutional Animal Care and Use Committee.

### Mouse cold housing

Mice were housed under 4°C for 24 to 48 hours. Regular 12h/12h light/dark cycles, unrestricted food and water were provided ad libitum.

### Monocyte fate mapping

CCR2^creER^ R26^TdTomato^ mice received a single oral gavage of tamoxifen (200 µL of 20 mg/mL tamoxifen in corn oil; MilliporeSigma #T5648, #C8267) to induce labeling of CCR2-expressing monocytes. On the same day, experimental mice were transferred to a 4 °C cold room for 24 hours, while control littermates were maintained at room temperature. Peripheral blood was collected to assess labeling efficiency using flow cytometry, and the proportion of TdTomato adrenal gland macrophages was compared against total adrenal macrophage population. For the fate mapping assessing the role of TGFβ signaling in monocyte recruitment, mice first received Ly573636 treatment 24 hours prior to cold exposure and TAM administration. While mice were in the cold room, Ly573636 was given again to ensure consistent TGFβ antagonism. Mice were sacrificed after another 24 hours of cold exposure. Flow cytometry was used to further analyze TdTomato+ macrophages and monocytes.

### Ly573636 treatment

The TGFβR inhibitor Ly573636 (Tasisulam; AdooQ #A13070) was reconstituted in DMSO at a concentration of 20 mg/mL and subsequently diluted to 5% DMSO in PBS. Experimental mice received 150 µL of diluted Ly573636 via intraperitoneal injection one day prior to cold exposure and daily throughout the 48-hour cold challenge. Control littermates received vehicle (5% DMSO in PBS) administered on the same schedule.

### Single cell-RNA-seq analysis

Publicly available scRNAseq covering acute cold stress (GSE161751^35^, GSE268521^29^), chronic atherosclerotic stress (GSE161751^35^, GSE268521^29^), chronic social defeat (GSE161751^35^) and acute *Candida albicans* infection (CRA007150^36^) were collected. All datasets were available in 10X structure except CRA007150. CRA007150 was downloaded as fastq files and turned into read counts using cell ranger count function. Data was imported in R using the Read10X function from Seurat (v4.2.1). Cells with fewer than 300 detected features were excluded as likely empty droplets, and genes with fewer than 10 total reads were removed to reduce sparsity. For quality control, we computed the per-cell mitochondrial RNA fraction and discarded cells with >25% mitochondrial content; we also calculated a ribosomal gene ratio and excluded ribosomal genes that showed strong skewing effects. Data were then normalized and scaled with NormalizeData and ScaleData, and variable features were identified with FindVariableFeatures. Datasets from cold challenge, atherosclerotic burden, and our previously published steady-state adrenal gland cohort were merged, and batch effects were corrected with RunHarmony. The Harmony embeddings were used to generate UMAP and t-SNE projections, using the RunUMAP, RunTSNE functions. The number of principal components was chosen by elbow plot, and clustering was performed with FindNeighbors and FindClusters. Cell types were annotated using SingleR, and clusters were renamed accordingly.

### Cell communication analysis

Cell-cell signaling was inferred with NicheNet (nichenetr) on our Seurat (v4.2.1) object, using adrenal macrophages as senders and endothelial cells as receivers, analyzed per condition. Genes were called expressed if present in ≥10% of cells with average log-normalized expression >0.1. EC differentially expressed genes (stress vs control) were identified with Seurat’s FindMarkers (two-sided Wilcoxon, Benjamini–Hochberg FDR < 0.05, no logFC threshold). Mouse symbols were mapped to human orthologs (biomaRt) before computing ligand activity with predict_ligand_activities. Ligands were ranked by score, and top candidates were intersected with expressed cognate receptors using the NicheNet ligand-receptor table.

### Flow cytometry

Freshly harvested adrenal glands were carefully trimmed of surrounding fat and placed in 90 µL of ice-cold RPMI medium (MilliporeSigma, #R8758-1L). Tissues were minced using fine scissors, followed by the addition of 10 µL Liberase™ DH (2.5 mg/mL in ddH_2_O; Roche, #5401054001). Samples were incubated at 37 °C for 35 minutes on an orbital shaker to achieve enzymatic dissociation. To halt further digestion, 1 mL of ice-cold FACS buffer (HBSS supplemented with 5% FBS and 2 mM EDTA) was added. The cell suspension was then passed through a 100 µm nylon mesh (McMaster-Carr) and the tube was then washed three times with buffer. Dissociated adrenal cells were pelleted by centrifugation at 1450 rpm for 5 minutes, the supernatant was discarded, and the remaining cells were resuspended in fresh cold FACS buffer for subsequent flow cytometry staining.

### Immunofluorescence imaging

Adrenal glands were fixed in 4% paraformaldehyde in 1× PBS for 1 h, embedded in OCT, frozen at −20 °C overnight, and cryosectioned at 10 µm. Sections were brought to room temperature (20 min), washed twice in PBS (5 min each), then blocked with 2.5% BSA (ThermoFisher #37525) and permeabilized with Tween-20 (Sigma-Aldrich #9005-64-5) for 45 min, followed by two PBS washes. Primary antibodies diluted (1:250) in PBS were applied overnight at 4 °C; slides were washed three times in PBS and incubated with secondary antibodies (1:500) for 1.5 h at room temperature, then washed three more times and mounted in Fluoromount (SouthernBiotech). Images were acquired on a Leica SP8 inverted confocal microscope. Quantification was performed in R using the tiff package (v0.1-8; readTIFF), reporting mean pixel intensity as the fluorescence measure.

### Evans blue dye assay

100 g of Evans blue dye was dissolved in 10 mL 1X PBS. Mice received 150 µL Evans blue dye via intraperitoneal injection. 1 hour after injection, mice were sacrificed and harvested. Both adrenal glands were completely minced up in 100 µL PBS 1X. Supernatant was collected in 96-well plates. Blue dye intensity was measured using a plate reader.

### Statistics

Analyses were performed in GraphPad Prism or R. Two-group comparisons used two-tailed unpaired t tests, with a minimum sample size of 3 per group. Comparisons involving more than two groups used ANOVA. P-values were adjusted for multiple testing with the Benjamini-Hochberg method unless noted. Adjusted P < 0.05 was considered significant. ns, *, **, *** and **** indicate not significant, P > 0.05, P ≤ 0.05, P ≤ 0.001 and P ≤ 0.0001 respectively.

## Data availability

All data will be made available upon request to the corresponding author.

## Acknowledgements

Research was supported by the National Institutes of Health (NIH) grants R01 AI165553 (J.W.W.), R01 HL166843 (J.W.W.), and MN Partnership for Biotechnology and Medical Genomics (J.W.W.). Additional support included American Heart Association (AHA) fellowship 903380 (M.T.P.), AHA 25PRE1361476 (H.H.), NIH F31 HL182131 (H.H.), NIH T32 HL166142 (M.D.C. and C.R.), and AHA 26POST1543363 (G.M.S.).

## Supplemental Figure Legends

**Sup Fig 1. Flow cytometry gating of adrenal gland macrophages**.

A–D. Flow cytometry gating scheme for blood monocytes in *Ccr2*^*GFP/WT*^ and *Ccr2*^*GFP/GFP*^ knock-out mice. E. Gating strategy for adrenal gland macrophages in *Ccr2-GFP* reporter mice. F–G. Quantification of adrenal gland macrophages. H–I. Quantification of GFP^+^ macrophages within MHCII-low and MHCII-high subsets.

**Sup Fig 2. Flow cytometry gating and quantification of adrenal gland macrophages**.

A. Flow cytometry gating strategy for adrenal macrophages and endothelial cells. B–C. Quantification of adrenal macrophages.

**Sup Fig 3. Flow cytometry gating of monocyte fate mapping**.

A. Flow cytometry gating scheme for adrenal monocyte fate mapping experiment, targeting TdTomato+ macrophages. B. Flow cytometry gating scheme for blood monocytes from the monocyte fate mapping experiment. C–L. Representative flow plots showing CCR2-TdTomato labeling efficiency and specificity, in control or Ly573636 treated mice.

**Sup Fig 4. Immunofluorescent imaging confirming adrenal gland macrophage TGFβ1 deletion**.

A. Validation of TGFβ1 deletion using IF. B. SMAD2 activity in TGFβ1-deficient mice indicated by pSMAD2 staining (green).

**Sup Fig 5. Morphological changes associated with adrenal gland macrophage TGFβ1 deletion**.

A–B. CD31^+^ endothelium morphology in TGFβ1-deficient or control mice.

**Sup Fig 6. Macrophage TGFβ1 deletion leads to elevated adrenal gland corticosterone**.

A. Measurement of adrenal gland corticosterone concentration from TGFβ1-deficient or control mice.

