## Supplemental Figures 1-6 for "Adrenal Gland Macrophage-derived TGF-β Governs Vascular Permeability to Drive Monocyte Recruitment during Stress"

### SUPPLEMENTAL FIGURE 1

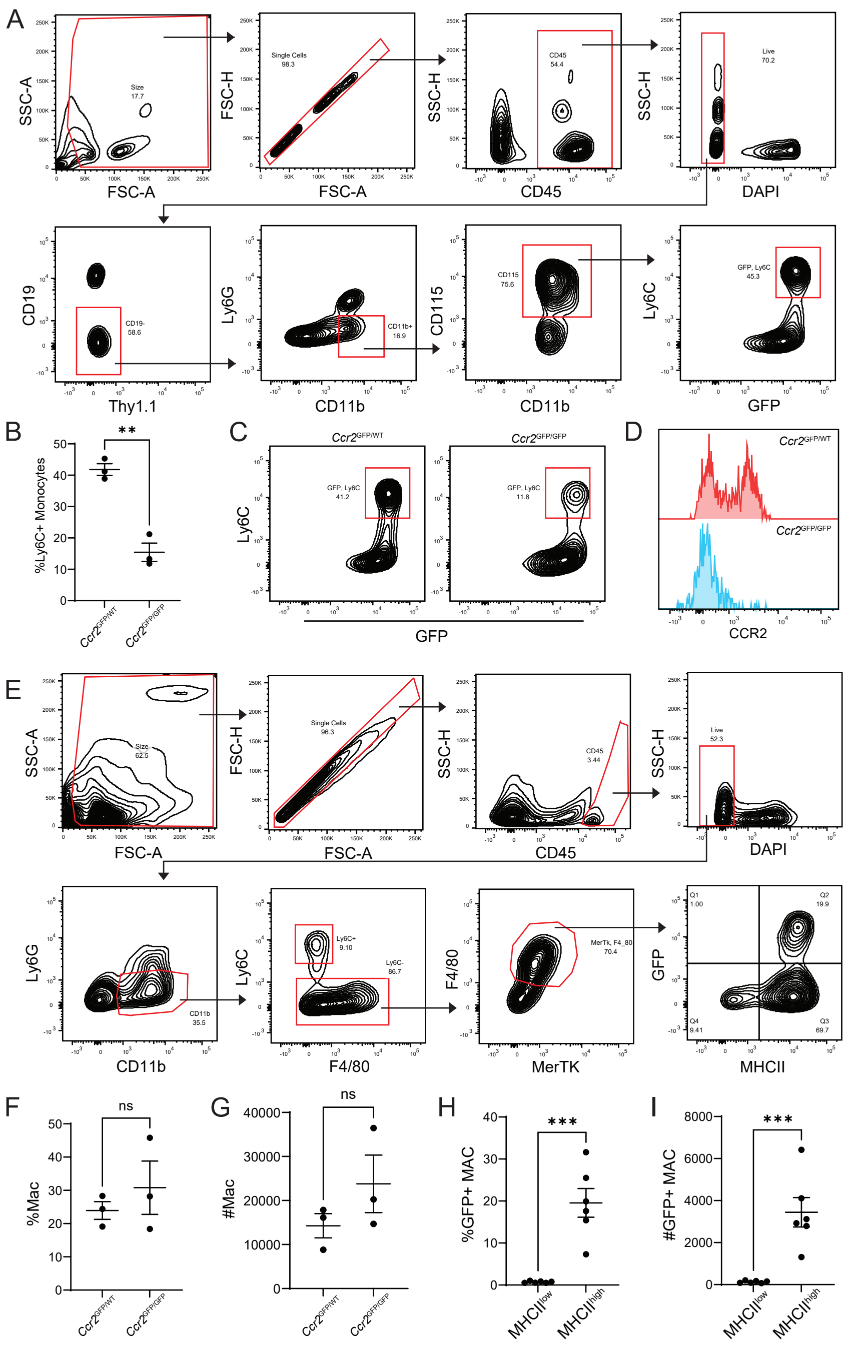

### SUPPLEMENTAL FIGURE 2

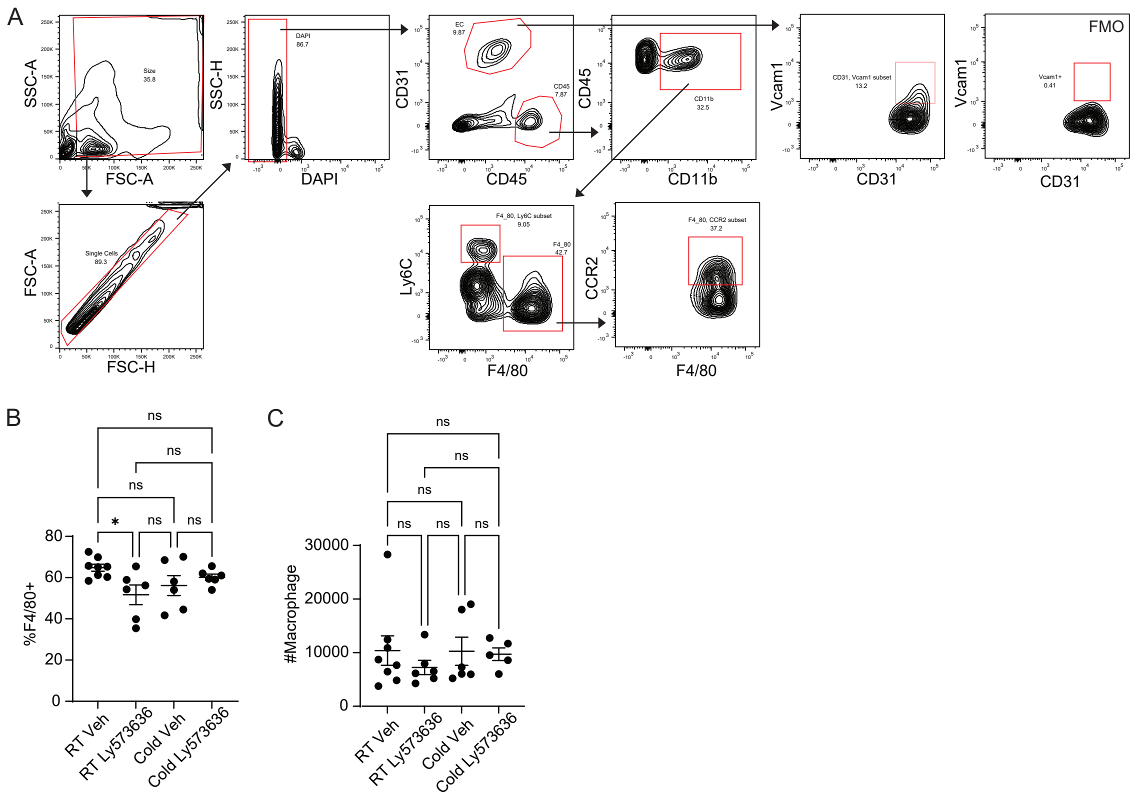

### SUPPLEMENTAL FIGURE 3

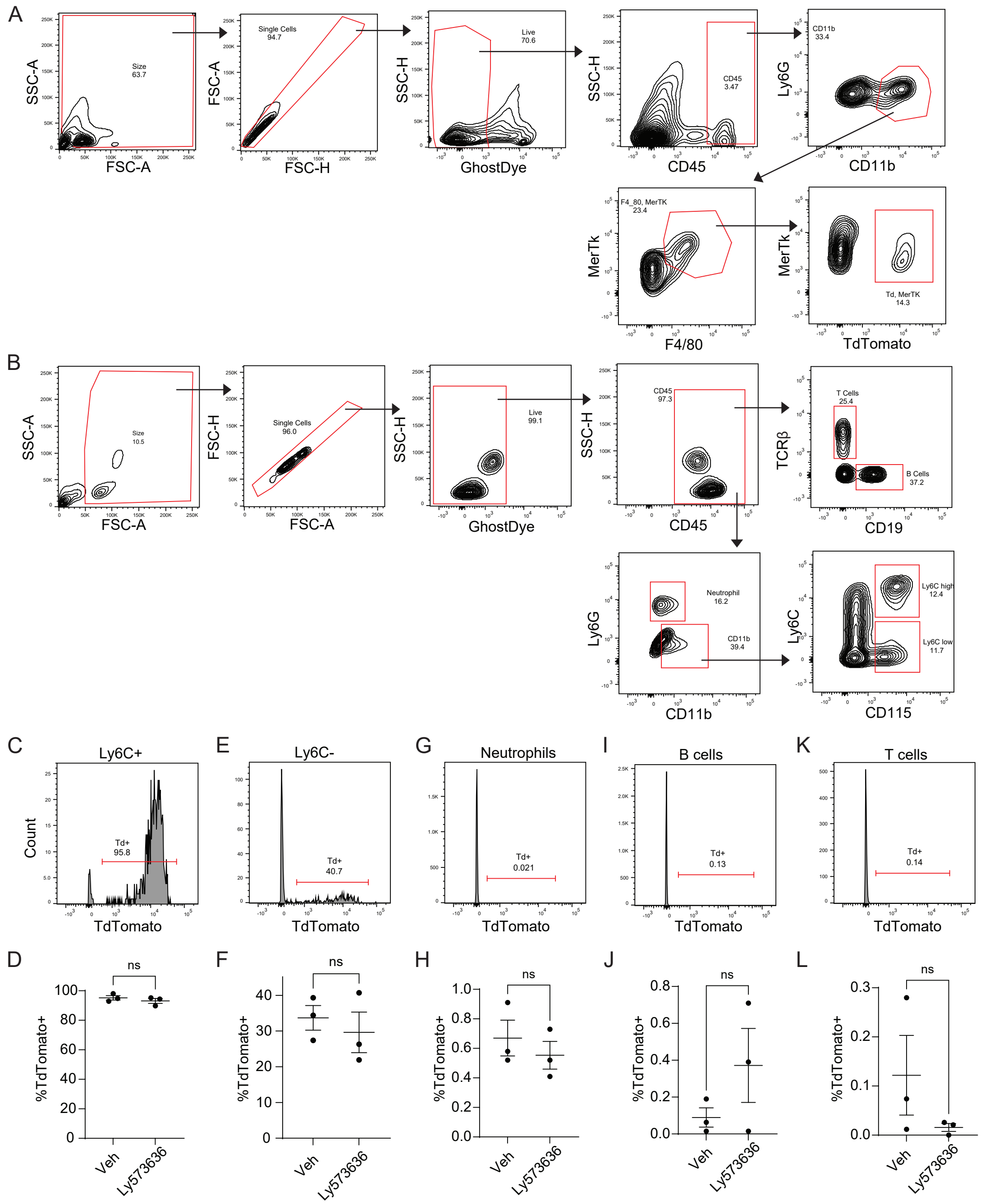

### SUPPLEMENTAL FIGURE 4

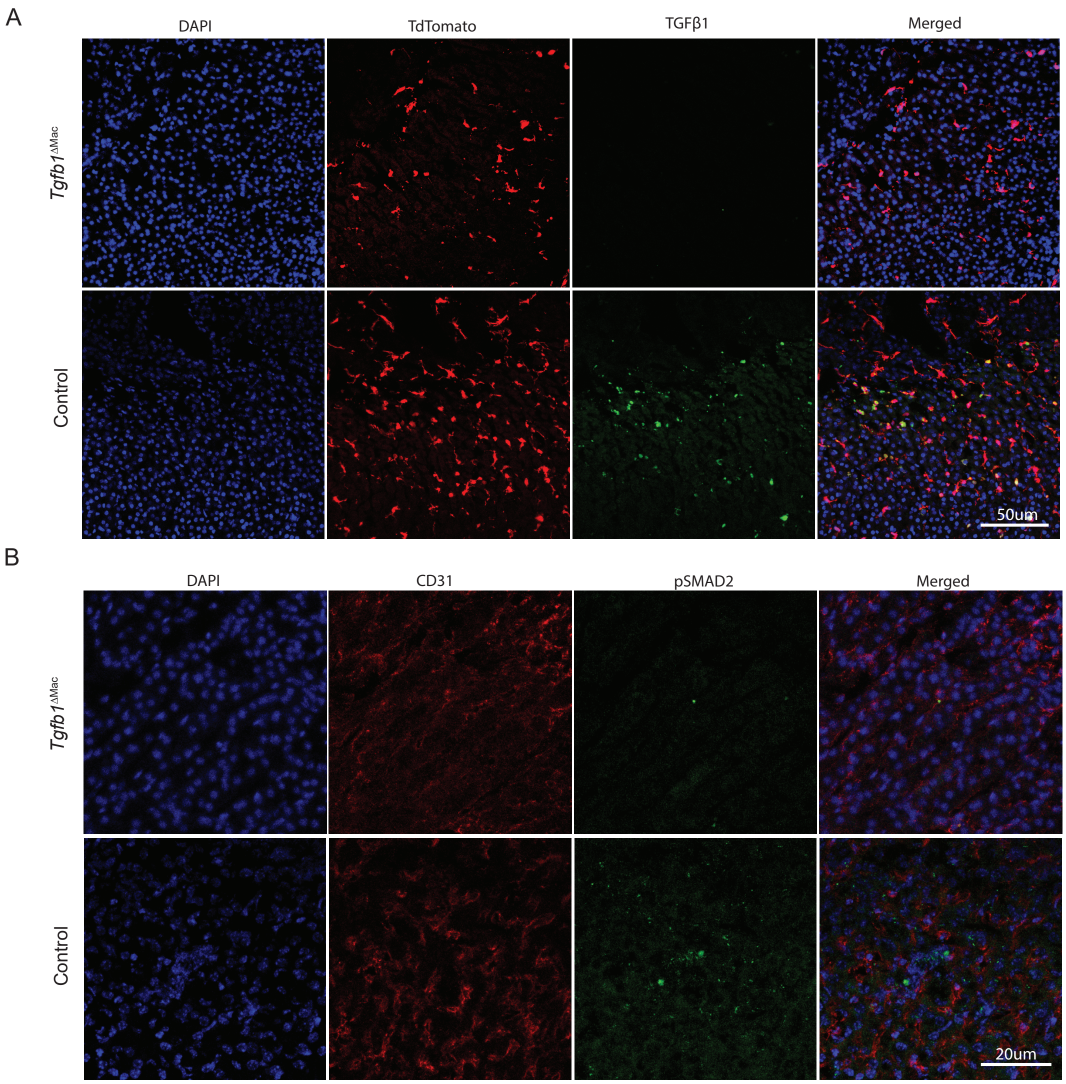

### SUPPLEMENTAL FIGURE 5

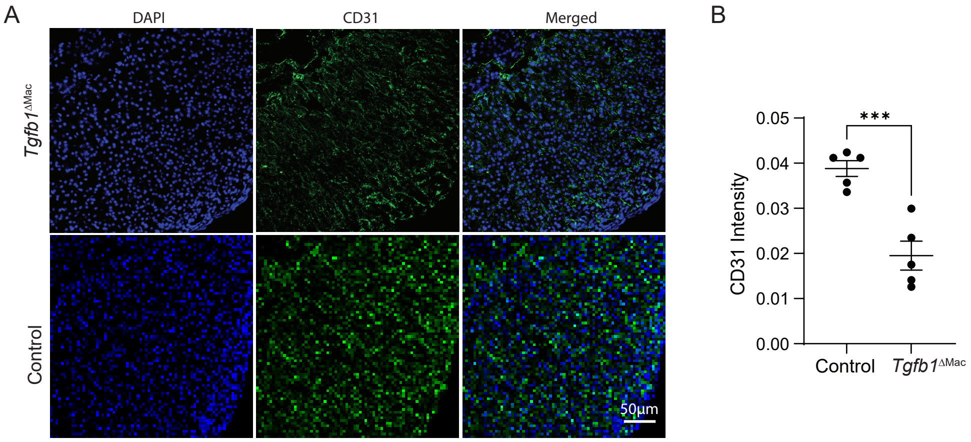

SUPPLEMENTAL FIGURE 6

A

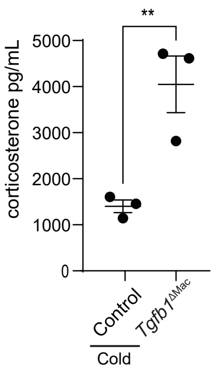
